# Drug-Mediated Control of Receptor Valency Enhances Immune Cell Potency

**DOI:** 10.1101/2023.01.04.522664

**Authors:** Paul B. Finn, Michael Chavez, Xinyi Chen, Haifeng Wang, Draven A. Rane, Jitendra Gurjar, Lei S. Qi

## Abstract

Designer T cells offer a novel paradigm for treating diseases like cancer, yet they are often hindered by target recognition evasion and limited in vivo control. To overcome these challenges, we develop valency-controlled receptors (VCRs), a novel class of synthetic receptors engineered to enable precise modulation of immune cell activity. VCRs use custom-designed valency-control ligands (VCLs) to modulate T cell signaling via spatial molecular clustering. Using multivalent DNA origami as VCL, we first establish that valency is important for tuning the activity of CD3-mediated immune activation. We then generate multivalent formats of clinically relevant drugs as VCL and incorporate VCR into the architecture of chimeric antigen receptors (CARs). Our data demonstrate that VCL-mediated VCRs can significantly amplify CAR activities and improve suboptimal CARs. Finally, through medicinal chemistry, we synthesize programmable, bioavailable VCL drugs that potentiate targeted immune response against low-antigen tumors both in vitro and in vivo. Our findings establish receptor valency as a core mechanism for enhancing CAR functionality and offer a synthetic chemical biology platform for strengthening customizable, potent, and safer cell therapies.

## INTRODUCTION

Immune cells make decisions by integrating a multitude of stimulatory and inhibitory signals via membrane-bound receptors^1,2^. Synthetic receptors tip the balance of these signals to generate favorable user-defined immunological outcomes, which has become a foundation for cell engineering and therapies for treating diseases such as cancer and autoimmunity^3–5^. Despite their promise, immune cells engineered with synthetic receptors are often hampered by issues including poor efficacy, target recognition evasion, and limited in vivo control^5–7^.

As one of the most clinically used synthetic receptors, chimeric antigen receptors (CARs) comprise single-chain transmembrane molecules that link user-defined tumor surface antigens to T cell receptor (TCR)-like signaling^6,8^. CAR T cell therapies targeting CD19 or BCMA have seen major clinical successes in treating blood malignancies^9–13^, yet several challenges remain in treating a wide range of tumors^14–17^. First, many CARs fail to exhibit robust in vivo activity and often require extensive protein engineering before achieving desired anti-tumor efficacy^18,19^. Second, engineered T cells frequently suffer from target recognition evasion, leading to antigen escape and suboptimal efficacy amid high tumor heterogeneity^20–22^. As examples, a reduction in antigen expression has been linked to cancer relapse in patients with B cell acute lymphoblastic leukemia (B-ALL)^23^. Furthermore, variable antigen densities of the heterogeneous cancer population greatly limit cell therapy efficacy in solid tumors^15,24^.

The ability to modulate *in vivo* T cell activity presents a promising strategy to overcome these limitations. It not only provides a versatile method to enhance the efficacy of poorly performing T cells, but also enables the same engineered cells to adaptively target tumors with variable and changing antigen densities^25^. If mediated by clinical drugs, this precise in vivo control may also offer a safety mechanism to mitigate adverse effects such as cytokine release syndrome and neurotoxicity^26–28^.

To enable modular and tunable control of T cell activity *in vivo*, we closely examined the biophysics of T cell receptor (TCR) signaling and compared it with that of CARs. TCR signaling operates on two dimensions, ‘vertical’ signaling between TCR and the antigen peptide-loaded Major Histocompatibility Complex (pMHC) and ‘horizontal’ signaling from spatial microclustering (**Fig. 1a**)^29^. While vertical binding events trigger mechanotransduction and subsequent kinase cascade activation^30^, the horizontal TCR clustering at the immunological synapse rapidly amplifies signaling, offering a potent mechanism for enhancing T cell cytotoxicity^31–36^. The degree of clustering (i.e., the number of molecules clustered) correlates with T cell potency and cytotoxicity^37–40^. In contrast, current CAR designs focus solely on cell-to-cell vertical signaling, mediated by antigen interaction with single-chain variable fragment (scFv)^41^. This design not only falls short of capturing the robust TCR signaling process, resulting in weaker T cell responses when targeting tumors with low antigen expression, but also misses opportunities to incorporate drug-mediated control mechanisms that could fine-tune receptor activity in vivo^42–44^.

**Figure 1.**
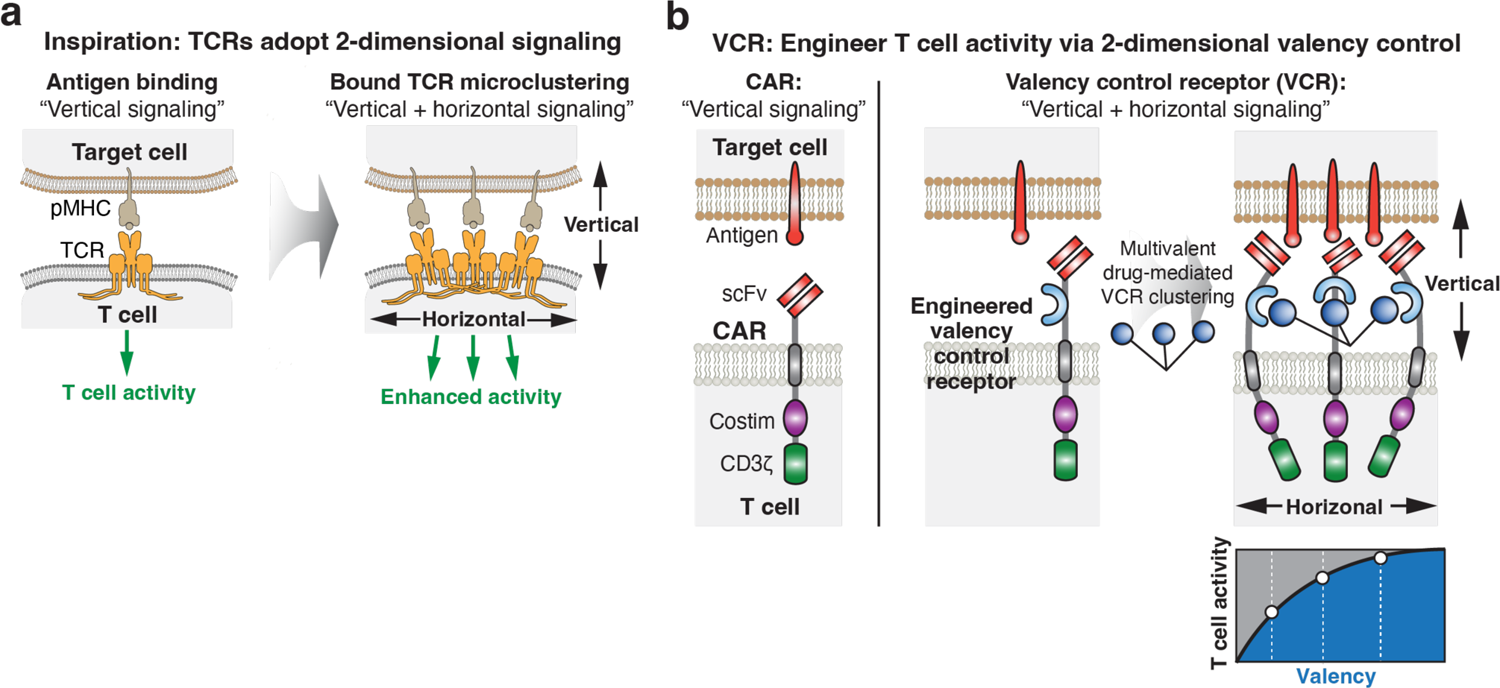
Concept of valency-controlled receptor (VCR) for drug-inducible enhancement of immune cell activity. **a**, TCR utilizes both ‘vertical’ signaling (binding between the extracellular domain and pMHC) and ‘horizontal’ signaling (microclustering) for potent T cell signaling. **b,** Schematic of the valency-controlled receptor (VCR) built upon the chimeric antigen receptor (CAR) molecule by combining vertical and horizontal signaling. VCR incorporates a multivalent drug binding domain into the extracellular hinge region to enable drug-mediated T cell activation. Dosing with a multivalent drug induces clustering of VCRs to enhance T cell signaling.

Here we developed valency-controlled receptors (VCRs), a novel class of synthetic receptors that can provide precise modulation of T cell activity via custom-designed valency-control ligands (VCLs) (**Fig. 1b**). By incorporating a drug-binding domain into the extracellular region of the receptor, we engineered a modular architecture that enables drug-inducible spatial receptor clustering. Utilizing multivalent DNA origami as ligands, we established the principle that valency control is a crucial aspect of receptor functionality and T cell activity. We incorporated the VCR domain into the architecture of chimeric antigen receptors (CARs) and demonstrated that VCR can significantly amplify CAR activity and even enhance suboptimal CARs. Through medicinal chemistry, we synthesized programmable, bioavailable, clinically relevant VCL drugs that potentiated targeted immune response against low-antigen tumors in vitro and in vivo. The VCR platform provides a new synthetic receptor platform for studying and controlling cell therapy via customizable drugs.

## RESULTS

### Developing a multivalent DNA origami ligand-mediated VCR system

Developing multivalent ligand-inducible VCRs necessitates engineering of both the ligand molecule and the receptor. Apart from the ligand-receptor binding affinity, multiple parameters can influence the receptor signaling, including receptor proximity, ligand valency, and microclustering dynamics. However, unlike DNA or protein engineering, medicinal chemistry can often be a slow and laborious process, limiting our ability to rapidly explore a large parameter space in parallel. To overcome this limitation, we first developed a flexible, facile, and versatile platform to characterize both the ligand molecule and the receptor architecture in parallel.

To establish a flexible VCL-VCR platform, we first adopted a receptor design that consisted of a single-stranded DNA (ssDNA) that was covalently conjugated via benzyl guanidine to an extracellular SNAP-tag protein (Fig. 2a)^39^. The SNAP-tag was fused to the intracellular signaling domains (CD28 and CD3ζ) and was connected by an IgG4 hinge and a CD28 transmembrane domain. This SNAP VCR (without scFv) can interact with a ssDNA ligand that is complementary to its conjugated ssDNA and their interaction can be programmed predictably using Watson-Crick base pairing. To precisely control VCR clustering, we developed a rigid Y-shape DNA origami scaffold that presented multiple ssDNA strands at the distal ends, which were complementary to ssDNA conjugated to the SNAP VCR (valency=3). Upon expressing SNAP VCR in Jurkat T lymphocyte cells, we found that VCL-mediated microclustering of VCRs can induce a modest level of T cell activation (Fig. 2b).

**Figure 2.**
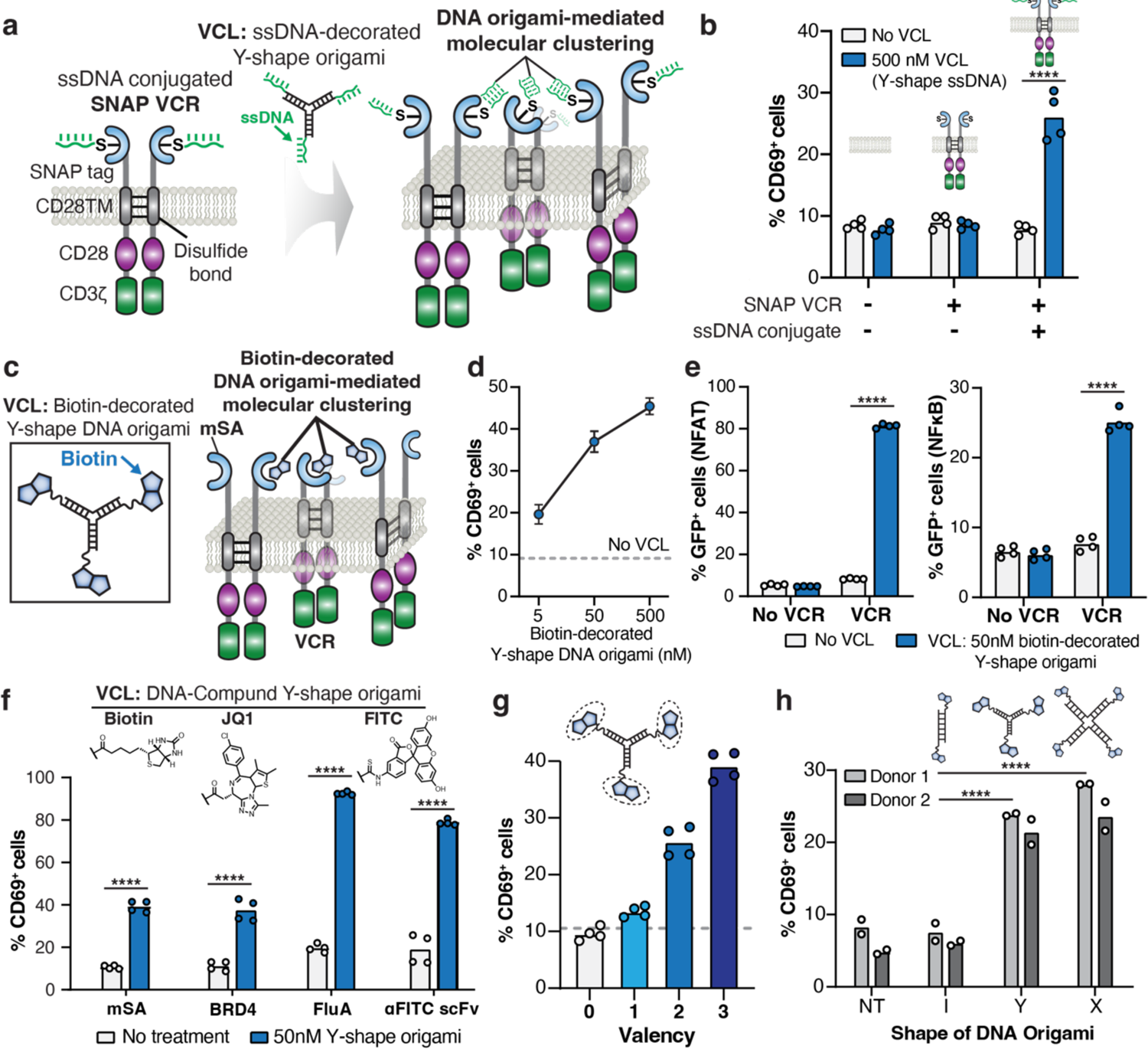
Development of a multivalent DNA origami-mediated VCR system and characterization of VCL-VCR interaction for T cell signaling. **a**, Development of a multivalent DNA origami-inducible VCR, SNAP-VCR, which contains an extracellular SNAP-tag protein, a CD28 transmembrane domain (CD28TM), a CD28 cytoplasmic domain, and a CD3ζ cytoplasmic domain. SNAP VCR is conjugated with a benzyl guanidine-modified single-stranded DNA (ssDNA) oligo. The DNA origami scaffold is utilized to create a trivalent ssDNA VCL (ssDNA Y-shape DNA) that induces clustering of the ssDNA-conjugated SNAP-VCR. **b,** SNAP-VCR induces Jurkat T cell activation when both the ssDNA-conjugated SNAP-VCR receptor and the ssDNA Y-DNA origami ligand (500 nM) are present. **c,** The Y-shape DNA origami is used to generate a trivalent biotin structure that interacts with VCR containing an extracellular monomeric streptavidin (mSA). **d,** mSA VCR Jurkat cells exhibit dose-dependent activation upon addition of the trivalent biotin Y-shape DNA origami VCL. **e,** mSA VCR induces NFAT (left) and NFκB signaling (right) in Jurkat reporter cells in the presence of trivalent biotin Y-shape DNA origami VCL. **f,** The Y-shape DNA origami is leveraged to create trivalent JQ1 and FITC and induce VCR Jurkat cells. The corresponding VCRs contain an extracellular BRD4 (for JQ1 Y-shape DNA), an FluA anticalin (for FITC Y-shape DNA), or a FITC scFv (for FITC Y-shape DNA). **g,** Y-shape DNA origami VCL can be linked to different numbers of biotin at its termini and used to probe the relationship between valency and T cell activation, which shows a positive correlation. **h,** Activation of two donors of primary human T cells using Y-shape or X-shape VCL but not for I-shape DNA. Data contain technical duplicates of at least two biological replicates or two donors and are statistically analyzed using students t test, **** = p < 0.0001.

We next sought to develop a more translatable system for using DNA origami to present small molecule moieties. To achieve this, we replaced the SNAP-tag domain on VCR with a monomeric streptavidin (mSA) domain^45^ and decorated the Y-shape DNA origami with covalently attached vitamin B7 (biotin) (Fig. 2c). Our results showed that Jurkat cells expressing the mSA VCR were activated by treatment with the biotin-decorated Y-DNA origami (valency=3) in a dose-dependent manner (Fig. 2d) and this activation occurred only when both VCR and VCL were present (Fig. 2e).

### Leveraging the programmability of DNA origami to characterize valency control

We next harnessed the rapid, facile, and programmable features of DNA origami design to characterize how VCL design affected VCR-mediated T cell signaling. We first asked whether different small molecules could be used to induce VCR activity. To address this, we swapped the mSA domain for the JQ1-binding human bromodomain-containing protein 4 (BRD4) domain, fluorescein isothiocyanate (FITC)-binding anticalin (FluA) protein domain, or a FITC-binding scFv. Correspondingly, we conjugated the Y-shape DNA origami with JQ1 or FITC (Fig. 2f). Our data showed that all designs induced T cell activation in the presence of the corresponding VCL, and that some ligand-VCR pairs induced stronger activation than others. This demonstrates that the versatility of VCRs, which can be controlled by different drugs in their multivalent format.

To determine how valency affected T cell activation, we synthesized a series of Y-shape DNA origami scaffolds decorated with different numbers of biotin (valency=0 to 3). The mSA VCR Jurkat cells induced by these ligands showed a positive correlation between T cell potency and valency (Fig. 2g). We further characterized valency parameters using different DNA origami scaffolds, including I-shape (valency=2), Y-shape (valency=3), and X-shape (valency=4), all of which were conjugated with biotin at the termini (Fig. 2h, **Fig. S1a**). Across three orders of VCL concentration, we observed consistently higher Jurkat T cell activation when using higher-valency VCLs (**Fig. S1b-c**). We next tested these design principles in primary human T cells. Two different donors expressing mSA VCR, both Y- and X-shape VCLs outperformed I-shape VCL (Fig. 2h). These experiments suggested that VCLs with valency greater than 2 were more effective in mediating VCR activity and that higher valency positively correlated with higher T cell activity.

Using DNA origami, we also assessed how the physical proximity of VCL-bound VCRs affected T cell activation. To address this, we increased the arm length of the Y-shape DNA origami by extending the number of nucleotides from 24 to 36 bp (**Fig. S2a**). Interestingly, the smaller VCL scaffold resulted in better T cell activation (**Fig. S2b**), which was also confirmed when examining activation-mediated IL-2 production (**Fig. S2c**). Collectively, our data imply that VCRs with closer proximity may lead to better T cell signaling.

### Development of multivalent small molecule VCLs to control VCRs

To generate a translatable VCL, we next sought to convert the multivalent DNA origami ligands into small molecule ligands. To achieve this, we utilized dendrimer scaffold chemistry. While 3- and 4-valency designs performed similarly in our DNA origami VCL system (Fig. 2h), the 4-valency design was more readily accessible through commercially available chemical scaffolds. We used peptide coupling chemistry to covalently conjugate biotin to a generation 0 PAMAM dendrimer, thereby generating a multivalent small molecule, termed V4 Biotin VCL (Fig. 3a). We then verified that V4 Biotin VCL could effectively activate the NFAT pathway in mSA VCR Jurkat reporter cells (Fig. 3b).

**Figure 3.**
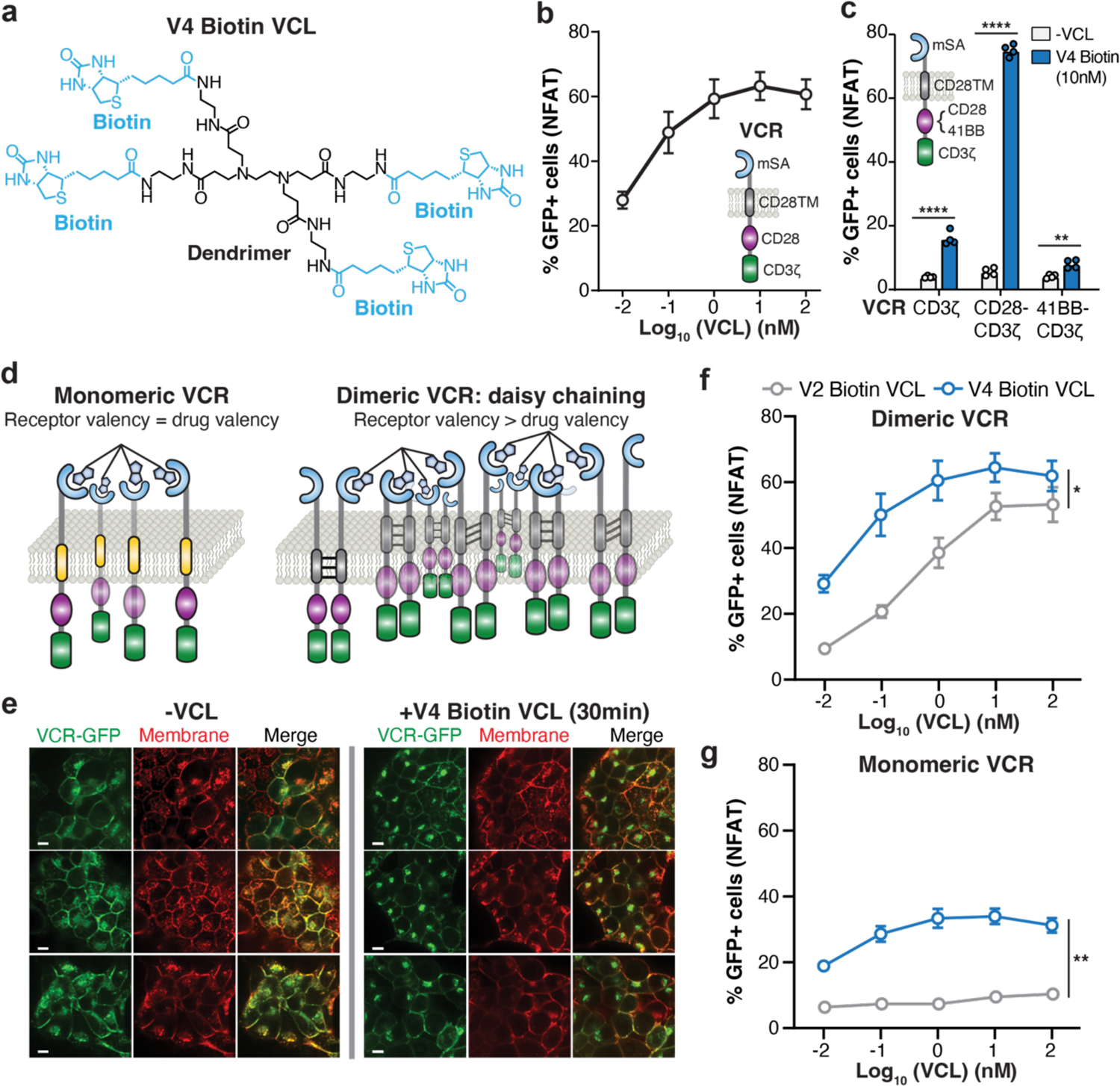
Development of multivalent small molecule VCL-inducible VCRs to modulate T cell activation and characterization of VCR clustering. **a**, Structure of the tetravalent biotin VCL synthesized using dendrimer chemistry (V4 Biotin VCL). **b,** V4 Biotin VCL-inducible VCR clustering triggers NFAT signaling in Jurkat reporter cells. **c,** Comparison of VCRs containing no, a CD28, or a 4-1BB costimulatory domain, with and without VCL, for NFAT signaling activation. **d,** Schematic showing how dimeric VCRs can lead to super-clustering of VCRs via ‘daisy chaining’, while monomeric VCRs presumably generate clusters with size that dictates the drug valency. **e,** Confocal fluorescent microscopy images of HEK293T cells expressing mSA VCR fused to GFP, with and without tetravalent V4 Biotin VCL for 30 minutes. Green, mSA VCR-GFP fusion; red, cell membrane stained with wheat germ agglutinin (WGA) conjugates CF568. **f-g,** Comparison of dimeric and monomeric VCR for triggering NFAT signaling activation, using tetravalent or bivalent biotin VCLs. Data contain technical duplicates of at least two biological replicates and are statistically analyzed using an unpaired student’s t test for **c** and paired for **f** and **g**, * = p < 0.05, ** = p < 0.01, **** = p < 0.0001.

Clinical CAR designs use either CD28 or 4-1BB as costimulatory signaling domains. Using V4 Biotin VCL, we compared whether the VCR system behaved similarly when using CD28 or 4-1BB in Jurkat cells. We observed that while VCR containing CD28-CD3ζ induced strong NFAT signaling with V4 Biotin VCL, VCR containing 4-1BB-CD3ζ induced almost no NFAT signaling, which was similar to VCR-CD3ζ alone (Fig. 3c).

### Multivalent VCL mediates super-clustering of VCR and amplifies T cell signaling

The IgG4 hinge and CD28 transmembrane domains used in the VCR system contain cysteine residues and hydrophobic residues that can form S-S bonds and π-π stacking interactions, respectively. These interactions result in the formation of dimeric VCRs on the cell surface, suggesting the possibility that the VCL is not simply creating small clusters of VCRs based on ligand valency, but instead is inducing “daisy-chaining” of many VCRs into larger clusters (Fig. 3d).

To test this hypothesis, we expressed an mSA VCR fused to a fluorescent protein (mSA-VCR-GFP) in HEK293T cells and performed confocal microscopy to visualize the formation of receptor clusters upon treatment with V4 Biotin VCL. After 30 minutes of VCL treatment, diffuse GFP signals formed large puncta at the cell surface, indicating the formation of VCR super-clusters (Fig. 3e).

We next developed a bivalent VCL (V2 Biotin VCL) (**Fig. S3**) and observed that this bivalent VCL also induced strong T cell activation using the dimeric VCR system, albeit to a lesser extent compared to V4 Biotin VCL (Fig. 3f). We further engineered a monomeric version of mSA VCR by mutating the cysteine residues (C47A and C78A) in the hinge domain and the hydrophobic residues (Y89L and T109L) in the transmembrane domain (**Fig. S4**). Interestingly, the monomeric VCR showed much weaker T cell activation when treated with V4 Biotin VCL and almost no T cell activation when treated with the bivalent VCL (Fig. 3g).

We further investigated if T cell signaling occurred due to VCR interactions on the same cell (cis) or from different cells (trans). We reasoned that if the effect is due to trans-interacting VCRs, increasing the cell density would also increase T cell signaling due to a higher likelihood for VCRs to interact (**Fig. S5a**). Notably, when cell densities were increased from 300 to 300,000 cells/cm^2^, there was no observable difference in T cell activation (**Fig. S5b**), indicating that VCL-induced signaling was largely due to clustering of VCRs on the same cell.

In summary, we conclude that VCR architectures containing cysteine residues and hydrophobic residues can form large super-clusters via a potential ‘daisy-chaining’ mechanism on the same cell, leading to greatly enhanced T cell signaling akin to TCR clustering.

### Development of the VCR-CAR fusion system for drug-inducible T cell activation

We incorporated the CD19 scFv (FMC63) onto the N-terminal end of mSA VCR and developed a VCR-CAR fusion system (Fig. 4a). We first compared the effect of 4-1BB and CD28 in this design. In the absence of CD19 antigen, VCR-CAR containing the CD28 domain exhibited strong NFAT activation when treated with V4 Biotin VCL, while VCR-CAR containing the 4-1BB domain retained low NFAT activation (Fig. 4b). This suggests that, while VCR-CD28 may exhibit antigen-independent CAR activity, VCR-4-1BB largely mitigates such effects. Therefore, we primarily used the 4-1BB costimulatory domain in subsequent studies.

**Figure 4.**
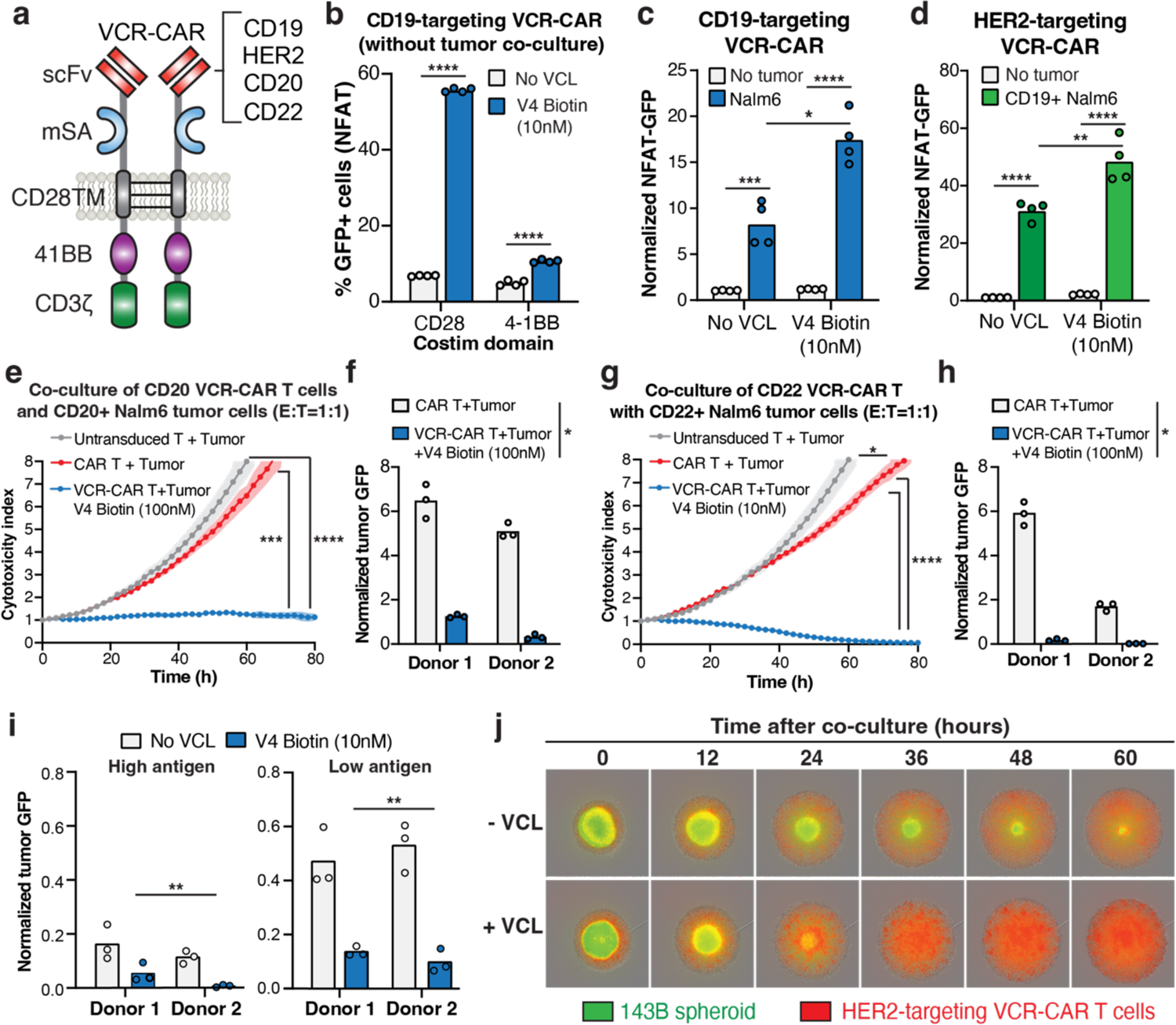
VCL-inducible VCR enhance CAR activity for better tumor killing. **a**, Schematic showing the VCR-CAR fusion molecule that allows using VCL to modulate the valency. Multiple CARs are tested including CD19, HER2, CD20, and CD22. **b,** Comparison of NFAT signaling with VCR-CAR containing a CD28 or 4-1BB costimulatory domain in the absence of tumor antigen, with and without 10nM V4 Biotin VCL. **c,** Characterization of CD19-targeting VCR-CAR (VCR-CD19 scFv-4-1BB-CD3ζ) for activating NFAT signaling in Jurkat cells with and without CD19^+^ Nalm6 tumor cell co-culture, and with and without 10nM V4 Biotin VCL treatment. **d,** Characterization of HER2-targeting VCR-CAR (VCR-HER2 scFv-4-1BB-CD3ζ) for activating NFAT signaling in Jurkat cells with and without CD19^+^ Nalm6 co-culture, and with and without 10nM V4 Biotin VCL treatment. Data for **b-d** show technical duplicates of two biological replicates and are statistically analyzed using a student’s t test, * = p < 0.05, ** = p < 0.01, *** = p < 0.001, **** = p < 0.0001. **e,** Real-time live-cell microscopy imaging of the co-culture of CD20-targeting VCR-CAR primary human T cells (VCR-CD20 scFv-4-1BB-CD3ζ) with CD20^+^ Nalm6 tumor cells (E:T ratio=1:1) in the presence of 100nM V4 Biotin VCL (blue), compared to untransduced T + tumor (grey) and CAR T + tumor co-culture (red) samples. **f,** Comparison of CD20-targeting VCR-CAR T (100nM V4 Biotin VCL) and CAR T at 60 hours post co-culture for reducing tumor cell densities across multiple donors. **g,** Real-time live-cell microscopy imaging of the co-culture of CD22-targeting VCR-CAR primary human T cells (VCR-CD22 scFv-4-1BB-CD3ζ) with CD22^+^ Nalm6 tumor cells (E:T ratio=1:1) in the presence of 10nM V4 Biotin VCL (blue), compared to untransduced T + tumor (grey) and CAR T + tumor co-culture (red) samples. **h,** Comparison of CD22-targeting VCR-CAR T (10nM V4 Biotin VCL) and CAR T at 60 hours post co-culture for reducing tumor cell densities across multiple donors. For **e** and **g**, data show means of technical triplicates and standard error from one donor and are statistically analyzed using a repeated measure ANOVA. For **e** and **g**, data show technical triplicates of two donors and are statistically analyzed using a paired student’s t test. * = p < 0.05, *** = p < 0.001, **** = p < 0.0001. **i,** Real-time live-cell microscopy characterization of tumor GFP signal using two donor T cells engineered with CD19-targeting VCR-CAR co-cultured with CD19_high_ (left) or CD19_low_ (right) Nalm6 tumor cells, with and without 10nM V4 Biotin VCL. **j,** Time-course microscopy images showing co-culture of HER2-targeting VCR-CAR primary T cells (red) and a low antigen spheroid model using HER2+ 143B bone osteosarcoma (green), without (top) and with (bottom) V4 Biotin VCL. Data for **f, h,** and **i** show technical triplicates of two donors and are statistically analyzed using a paired student’s t test, * = p < 0.05, ** = p < 0.01, *** = p < 0.001, **** = p < 0.0001.

To test whether VCR-mediated signaling synergizes with tumor antigen-induced T cell activation signaling, we co-cultured VCR-CAR Jurkat cells with CD19^+^ Nalm6 leukemia cells both in the presence and absence of V4 Biotin VCL (Fig. 4c). We found that VCR clustering amplified CD19-induced NFAT activity, leading to a significant enhancement in T cell activation when VCL was present. This amplification effect was not limited to CD19. We also observed enhanced T cell activation when targeting clinically relevant solid tumor antigens, including human epidermal growth factor receptor 2 (HER2). When co-cultured with HER2^+^ Nalm6 cells, the VCR-CAR system (HER2 scFv-mSA-4-1BB-CD3ζ) showed a similar VCR-mediated enhancement of T cell activity, underscoring the system’s modularity in relation to various tumor antigens (Fig. 4d).

### VCR enhances the CAR activity for better tumor killing

Engineered CARs often exhibit suboptimal anti-tumor activity and we wondered if appending the valency control domain to induce clustering can improve their efficacy. To do so, we fused the V4 Biotin VCL-controlled VCR domain onto CD20 CAR or CD22 CAR and compared the performance of VCR-CAR to CAR alone. While CD20 CAR and CD22 CAR showed suboptimal tumor killing activity, we observed significantly improved tumor killing activity using VCR-CAR in the presence of VCL (Fig. 4e-h). These findings were corroborated across multiple donors and various effector-to-target (E:T) ratios (**Fig. S6a-d**). These data raised the possibility that incorporating drug-mediated molecular clustering domains can modularly enhance the tumor-killing abilities of otherwise suboptimal CARs.

A common evasion strategy employed by tumors targeted by CAR T cells is downregulation of antigen expression. We therefore hypothesized that receptor clustering by our VCR platform could sensitize CAR T cells with low antigen density tumors. To evaluate this, we co-cultured VCR-CAR T cells with a recently developed CD19_low_ Nalm6 tumor model^25^. VCR-CAR T cells from different donors with VCL treatment vastly outperformed those without VCL treatment, indicating that low antigen tumor recognition can be improved by inducing a higher receptor valency (Fig. 4i).

We further verified the performance of VCR-CAR T cells in a low-antigen tumor spheroid model. We co-cultured HER2-targeting VCR-CAR (HER2 scFv-4-1BB-CD3ζ) T cells with HER2+ 143B osteosarcoma spheroids and noted greatly enhanced killing activity by VCR-CAR T cells when treated with VCL (Fig. 4j).

### Custom-designed VCL drugs enable effective VCR-CAR mediated tumor killing *in vivo*

To translate our VCR strategy into a viable therapeutic strategy, we designed and characterized a series of small molecule-based biotin VCLs by varying their linker design (VCL1∼4, Fig. 5a). We assessed their half-life *in vivo* and determined VCL2 and VCL4’s half-life around 3∼4 hours and VCL1 and VCL3 were less stable with a half-life of fewer than 15 minutes (Fig. 5b). Next, we quantified VCL-induced T cell activation using the NFAT Jurkat cells and found that VCL4 performed significantly better at low concentrations (pM to sub-nM) compared to VCL1 or VCL2 (Fig. 5c **& Fig. S7**). Based on these results, we selected VCL4 for further in vivo efficacy studies.

**Figure 5.**
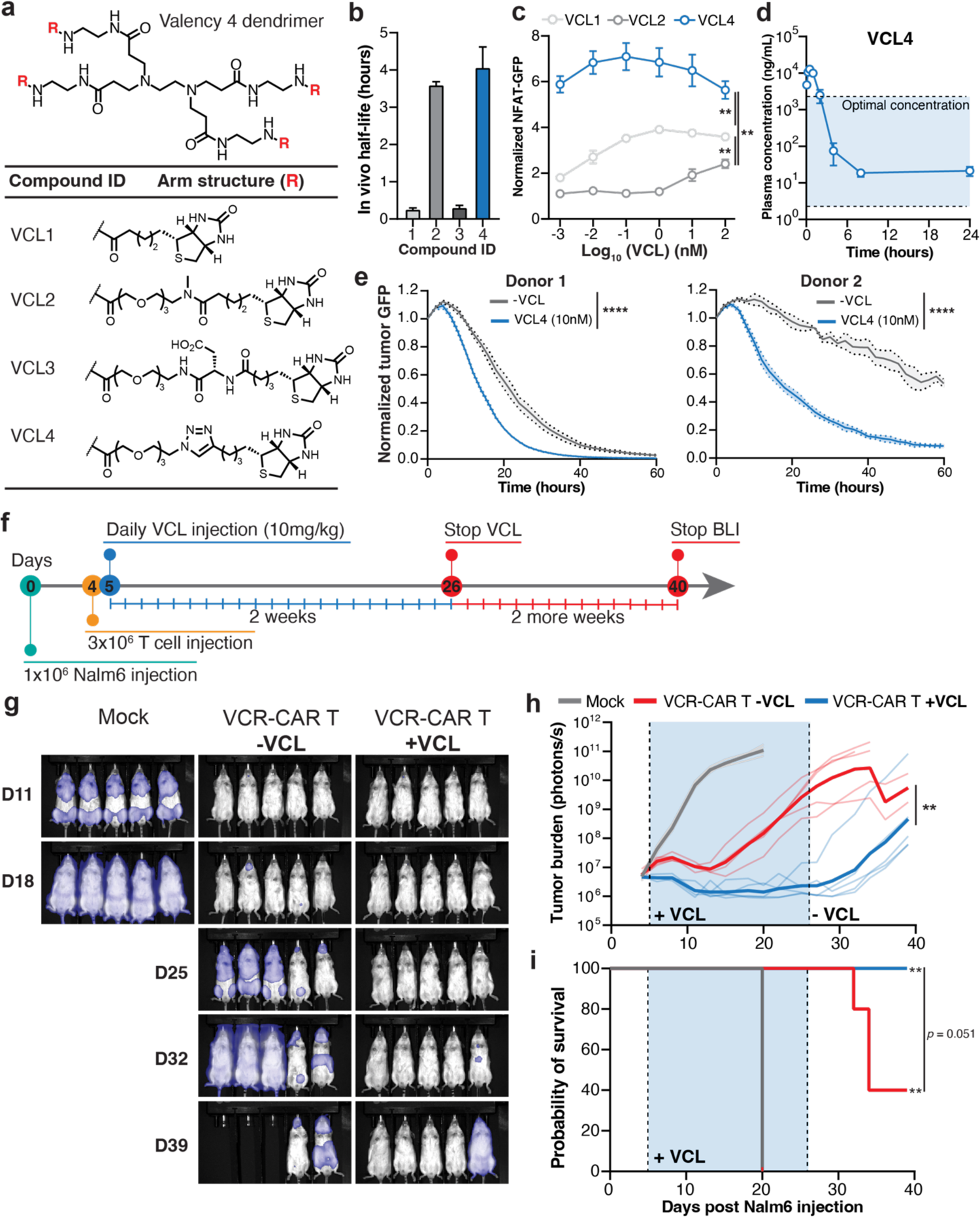
Development of custom-designed, multivalent, bioavailable VCL drug to modulate T cell activity in vivo. **a**, Design of four tetravalent biotin VCL drugs (VCL1-4) conjugated to a generation 0 PAMAM dendrimer with different linkers to improve bioavailability. **b,** Characterization of half-life of the four lead series compounds in mice blood plasma (n = 3). **c,** Characterization of VCL1, VCL2, and VCL4 for activating NFAT signaling in Jurkat cells using mSA VCR. Data show technical duplicates of two biological replicates and are statistically analyzed using paired student’s t test, ** = p < 0.01. **d,** Characterization of VCL4 plasma concentration in the mouse serum over 24 hours (n = 3). **e,** Characterization of tumor signal using real-time live cell microscopy for the co-culture of CD19-targeting VCR-CAR primary T cells and CD19+ Nalm6 cells for two donors, without (grey) and with (blue) 10nM VCL4 treatment. Data of each donor show mean and standard deviation of three technical replicates and are statistically analyzed using repeated measure ANOVA, **** = p < 0.0001 for both donors. **f,** Schematic overview of the animal experiment. 1×10^6^ CD19^+^ Nalm6 cells are i.v. injected into mice and 4 days later, 3×10^6^ T cells (mock) or 3×10^6^ VCR-CAR T cells (CD19 scFv-mSA-4-1BB-CD3ζ) are i.v. injected. One group of mice are dosed by i.p. injection with 10mg/kg VCL4 daily for 21 days and tumor burden is measured by BLI three times each week. **g,** BLI images showing groups of mice injected with mock T cells, VCR-CAR T cells without VCL, and VCR-CAR T cells with VCL. Light lines show individual mice while dark lines represent mean. Data are statistically analyzed using repeated measure ANOVA, ** = p < 0.01. **h,** Characterization of tumor burden (photon/s) for mice injected with VCR-CAR T cells without VCL (red) or with VCL (blue), and mock (grey). **i,** Survivial curve for VCL-dosed mice carrying VCR-CAR T cells without VCL (red) or with VCL (blue), and mock (grey). In **h-i**, data are analyzed using a log rank test compared to mock, ** p < 0.01, or comparing the two VCR groups (p = 0.051).

We then monitored the plasma concentration of VCL4 following a 10 mg/kg intraperitoneal injection (i.p.) in wildtype mice and found that the drug maintained an effective concentration over a 24-hour period (Fig. 5d). This implies that a daily drug dosing schedule of 10 mg/kg could adequately sustain in vivo VCR-CAR activation. VCL4 could synergize with antigen-mediated T cell signaling, amplifying the cytotoxicity of VCR-CAR T cells against CD19^+^ Nalm6 tumor cells in an in vitro setting using different donor T cells (Fig. 5e).

To assess the in vivo activity of VCR-CAR T cells, we injected 1×10^6^ luciferase-expressing CD19^+^ Nalm6 cells by tail vein injection into a NOD/SCID mouse, followed by 3×10^6^ VCR-CAR T cells tail vein injection four days later (Fig. 5f). Mice were treated with VCL4 at 10 mg/kg (i.p.) daily for a total of 21 days, and tumor burden was routinely measured by bioluminescence imaging (BLI) three times a week. Subsequent BLI measurements were conducted an additional 14 days post-VCL withdrawal to gauge the reversible effects of the therapy. Mice treated with a combination of VCR-CAR T cells and VCL4 showed a significant reduction in tumor burden compared to the control groups (Fig. 5g-h). Crucially, VCL4 treatment bestowed a strong survival advantage during its administration (Fig. 5i).

To further evaluate the in vivo effectiveness of VCR-CAR T cells against low-antigen tumors, we utilized a CD19_low_ Nalm6 mouse model, a challenging target for conventional CAR therapies^25^. Following the same dosing schedule (Fig. 5f), mice treated with VCR-CAR T cells and VCL4 manifested a marked decrease in tumor burden compared to controls (**Fig. S8a-b**). Importantly, VCL4 dosing conferred a significant survival advantage, while all mice without VCL performed only marginally better than mock and succumbed to the cancer (**Fig. S8c**). Collectively, our results indicate that VCR-CAR T cells offer a drug-responsive means of enhancing the targeting of low-antigen tumors.

## DISCUSSION

In this study, we developed a novel receptor platform, termed valency-controlled receptors (VCRs), that allows one to harness clinically relevant drugs to control synthetic receptor valency and modulate T cell activation and tumor killing. Our VCRs recapitulate TCR horizontal signaling, which has been implicated as an important mechanism for immune response regulation^46^. By precisely controlling the valency of receptors, we enhance the tunability, efficacy, and safety of engineered T cell therapies. With this method, our work further established the importance of valency control in fine-tuning the activity of costimulatory domain signaling and CAR activity, offering a new mechanism that can be explored for immunological engineering.

We designed different types of multivalent VCL ligands, including DNA origami, DNA origami-small molecule conjugate, small molecule ligands (biotin, JQ1, FITC), and custom-designed drugs, all of which have been utilized to modulate VCR activity. This demonstrates the high versatility and generalizability of the system. These ligands allowed us to probe the biophysical parameters influencing VCR performance. For example, by leveraging the easy programmability feature of DNA origami VCL, we characterized both receptor (i.e., cluster size and proximity) and ligand (i.e., valency and concentration) parameters for T cell activation, without the need for sophisticated medicinal chemistry. Additionally, our system’s designability empowered us to create a series of small molecule VCLs for dose-dependent control of VCR T cell activity, both in vitro and in vivo.

Suboptimal performance of CARs has long been a bottleneck in the field, typically requiring laborious protein engineering and validation. We propose that valency control could enable many suboptimal CARs to achieve better efficacy via controlled receptor clustering. Supporting this, we successfully integrated VCR into various CAR systems, including those targeting CD19, HER2, CD20, and CD22, demonstrating enhanced CAR T cell cytotoxicity upon VCL drug treatment. Notably, converting these CARs to the VCR format simply requires modular addition of a fragment (∼1kb), making it seamlessly compatible with existing cell manufacturing processes. This expedites the production pipeline and extends its utility across various CAR systems.

For cancer immunotherapy applications, the ability to precisely tune CAR activity *in vivo* during patient treatment using customized drugs has an important impact on clinical outcomes. Several recent drug-inducible systems have been designed, usually around a single drug molecule with difficult-to-tune pharmacokinetic properties^47–50^. In comparison, the VCR platform provides a unique solution for the rapid, programmable design of customized multivalent drugs. Since the ligand-receptor occurs outside of cells, unlike in other drug-inducible systems, VCL drugs do not need to enter cells and thus avoids the cell membrane penetration issue of many drug designs. Many clinically relevant drugs could be converted into a multivalent format as VCLs to modulate VCR activity *in vivo*, which expands the repertoire of small molecules that can be repurposed for safely controlling cell therapy *in vivo*.

By integrating customized small molecule drug control into T cell therapy, the VCR platform enables clinicians to make decisions on the timing and potency of the therapy, thus creating novel avenues for safer and more effective treatments for a myriad of cancers and other diseases. In a possible clinical setting using VCR T cells, the drug dosing schedule could be modified as tumor cells undergo reduction of antigen expression. Additionally, dosing schedules could be tested to limit potential side-effects and increase safety by reducing or stopping drug treatment. Moreover, tumor localized drugs (e.g., conjugated to an antibody that binds to tumor antigens) could be explored to ensure T cells are only activated at the site of cancer, reducing the off-target effects often seen in solid tumor CAR T cells^51,52^. As our understanding deepens on how to induce VCR T cells *in vivo*, more complex dosing schedules could be leveraged such as oscillatory regiments that could help overcome T cell dysfunction^53,54^. Together, the VCR platform creates a novel avenue to better control the response of broad CAR T cell applications.

## METHODS

### Generation of VCR constructs

Standard molecular cloning techniques were used to assemble constructs in this paper. The SNAP protein was amplified from the DNA-CARzeta-GFP plasmid (a gift from Ron Vale, Addgene # 89344). The monomeric streptavidin (mSA), bromodomain 4 (BRD4), IgG4 hinges, CD137, CD28, and CD3z sequences were cloned from ordered gBlocks (Integrated DNA Technologies). The anti-FITC scFv and anti-FITC anticalin (FluA) were cloned from synthesized DNA (Twist Biosciences). The VCR-19BBz and VCR-HER2BBz were cloned from plasmids containing 19BBz (Crystal Mackall, Stanford University) and HER2BBz (Robbie Majzner, Stanford University), respectively. Plasmids were cloned using InFusion (Takara Bio) and Stellar Competent cells (Takara Bio).

### Cell lines

Jurkat Clone E6-1 cells were obtained from ATCC (TIB-152). The EGFP NFkB reporter Jurkat cell line was a gift from X. Lin (Tsinghua University). The EGFP NFAT reporter Jurkat cell line was a gift from A. Weiss (University of California, San Francisco). The Jurkat cell lines, Nalm6 cell lines, and 143b osteosarcoma cells were cultured in RPMI 1640 (Thermo Fisher) supplemented with 10% fetal bovine serum (FBS) (Alstem) and 100 U mL^-1^ of penicillin and streptomycin (Gibco) (complete RPMI). LentiX HEK 293T cells (Clontech) were cultured in DMEM + GlutaMAX (Thermo Fisher) supplemented with 10% FBS and 100 U mL^-1^ Pen/Strep (complete DMEM). Jurkat cells and Nalm6 cells were passaged every 2-3 days as cell culture densities approached 1 x 10^6^ cells per mL and 2 x 10^6^ cells per mL, respectively. LentiX HEK293T and 143b osteosarcoma cells adherent cell lines were passaged every 2-3 days by dissociating in 0.05% trypsin (Life Technologies) for 5 min at 37 °C or 1x DPBS supplemented with 2% FBS and 5 mM EDTA for 10 - 15 min at 37 °C to remove from culture plates. Cells were maintained at 37 °C and 5% CO_2_ and passaged using standard cell culture techniques. Cells were not tested for mycoplasma contamination.

### Stable cell generation

Stable Jurkat cells lines were generated using lentiviral transduction. Jurkat cells were seeded the day of transduction at 1 x 10^5^ cells per mL in 500 uL complete RPMI in a 24 well plate. For lentiviral production (method 1, six-well plate format), 3 x 10^5^ LentiX HEK293T cells in 2 mL complete DMEM were transfected with 1.51 ug of pHR vector with VCR construct, 1.32 ug dR8.91 and 165 ng of pMD2g with 7.5 uL of Mirus TransIT-LT1 in 250 uL OptiMEM (Thermo Fisher). Three days post transfection, lentivirus was harvested and filtered through a 0.45 um polyvinylidene fluoride filter (Millipore). After filtration, lentivirus was mixed 4:1 with Lentivirus Precipitation Solution (Alstem) and refrigerated overnight. Lentivirus was pelleted at 1,500 x g for 30 min at 4 C and resuspended in complete RPMI. Cells were treated with 50 - 100 uL of resuspended lentivirus (method 1) for 18 - 24 h. After 24 h, fresh media was exchanged and transduced cells were cultured for 5 - 7 days before assay.

### Generation of VCR-expressing primary T cells

Mixed CD4+ and CD8+ primary T cells were isolated with the Dynabeads™ Human T-Activator CD3/CD28 for T Cell Expansion and Activation (Thermo Fisher) from healthy donor whole blood obtained from the Stanford Blood Center (Stanford, CA) and stored in LN2. Thawed T-cells were cultured in complete RPMI 1640 with 100 U/mL rIL-2 (Thermo Fisher). T cells were stimulated with CD3/CD28 Dynabeads (Thermo Fisher Scientific) at a 2:1 cell:bead ratio for 1 d were transduced with lentivirus (method 2). Dyna beads were removed after 3 d of culture and adjusted every 2 days to maintain a cell density of 1 x 10^6^ - 2 x 10^6^ cells per mL. All T cell activation, cytotoxicity, and cytokine in vitro assays were performed on day 11 after bead activation. For lentivirus production (method 2, six-well plate format), the day before transfection 1.5 x 10^6^ LentiX HEK293T cells were plated in 2 mL complete DMEM. The day of transfection, 1.0 mL of complete DMEM was removed and cells were transfected with 1.79 ug of pHR vector with VCR construct, 1.28 ug psPAX2 and 554 ng of pMD2g with 10.9 uL of Mirus TransIT-LT1 in 435 uL OptiMEM (Thermo Fisher). At 6 h post transfection, media was exchanged with fresh, complete DMEM containing 1x ViralBoost reagent (Alstem). At 24 h post transfection lentivirus was harvested, filtered, concentrated, and resuspended in complete RPMI.

### Synthesis of small molecule-conjugated DNA origami

DNA origami structures were prepared immediately before use by annealing complementary ssDNA strands at 100 uM using the following program: 95 °C for 5 min, 65 °C for 5 min, 60 °C for 5 min followed by slowly lowering temperature to 4 °C. DNA origami was stored at 4 °C until used.

### BG-DNA labeling of Jurkat cells expressing SNAP VCR

Jurkat cells expressing SNAP-VCR were pelleted then washed twice and resuspended in HEPES buffered saline (HBS: 20 mM HEPES - pH 7.4, 10 mM Glucose, 4 mM KCl, 135 mM NaCl, 0.9 mM CaCl_2_, 0.5 mM MgCl_2_) with 0.5% BSA (HBS-BSA). SNAP-VCR Jurkat cells were conjugated with or without ssDNA-BG by incubating 2.0 x 10^7^ cell/mL in 200 uL with 5 mM ssDNA-BG or blank HBS-BSA (control) at room temperature for 30 min. During conjugation cells were gently agitated every 5 min to keep suspended. DNA-conjugated Jurkat cells were washed twice and resuspended in assay media before being used.

### Flow cytometry of CD69 expression and NFAT/NFkB reporter assays

Jurkat cells expressing the indicated construct were seeded at 5 x 10^4^ cells per well in 96-well format, with or without DNA origami or biotin small molecule, total volume 100 uL per well. Cells were stained for CD69 (clone FN50, BioLegend, dilution 1:500 in PBS + 2% FBS) expression after 4 - 24 hr at 37 °C. Reporter induction was assessed by flow cytometry (Beckman-Coulter Cytoflex S) after 16 - 20 hr (NFAT EGFP) or 20 - 24 hr (NfKb) at 37 °C. For co-culture assays, 5 x 10^4^ target cells per well were also added (effector:target E:T ratio = 1:1), total volume 100 uL per well. Cells were analyzed in 96-well format and 10,000 cells from the population of interest were collected for each sample. Data were analyzed using FlowJo v.10.8.1 (BD Biosciences).

### Cytokine production quantification

Jurkat cells expressing the indicated construct were seeded at 2.5 x 10^5^ cells per 500 uL in 24-well format, with or without Biotin-conjugated DNA origami. Supernatants from cell cultures were harvested after 24 h and samples were stored at −80°C. Secreted proteins were measured using the ELISA MAX Deluxe kits (Biolegend). Absorbance was measured at 450 nm and 570 nm using a Synergy H1 plate reader (BioTek), and protein concentrations were calculated by standard curves fitted to a power law.

### Confocal microscopy

To visualize VCR membrane localization, HEK293T cells expressing a low level of mSA VCR-eGFP were imaged in living cells 30 minutes after adding the biotin VCL. Both treated cells and control cells were stained with wheat germ agglutinin (WGA) conjugates CF568 (Biotium 29077-1) to visualize the plasma membrane. Images were taken on a Nikon TiE inverted spinning disk confocal microscope (SDCM) equipped with a Photometrics Prime 95B camera, a CSU-X1 confocal scanner unit with microlenses, and 405 nm, 488 nm, 561 nm, and 642 nm lasers. Images were taken using NIS Elements version 4.60 software with Z stacks at 0.3 μm steps, using the 60 × PLAN APO IR water objective (NA = 1.27). Scale bar = 10um.

### T cell cytotoxicity assays

Cytotoxicity assays were performed by co-culturing 50,000 GFP+ tumor cell targets with engineered T cells at the indicated E:T ratios with or without biotin small molecule in 200 uL of complete RPMI 1640 (without IL2) in a 96-well plate. Wells were mixed well and cells were allowed to settle for 1-2 h. Four images per well were acquired at 10x magnification in the green channel (200 ms exposure) and the red channel (600 ms exposure) every 2 to 3 hrs using an Incucyte (Sartorius). A cytotoxicity index was calculated by dividing total green fluorescence intensity per mm^2^ at every time point by the intensity per mm^2^ at the first time point. For spheroid experiments, HER2+143b osteosarcoma cells were plated at 4 x 10^5^ cells per well in 100 uL in ultralow attachment U bottom 96-well plate and allowed to form spheroids for 36 - 48 h before T cells and 10nM V4 Biotin VCL were added without mixing. One image per well was acquired at 4x magnification in the green channel (100 ms exposure) and the red channel (600 ms exposure) every 2 to 3 hr using an Incucyte (Sartorius).

### Bioavailability of drug molecules

For each drug molecule tested, two groups of three male CD-1/C57BL/6 mice were injected by intraperitoneal (IP) injection at 10 mg/kg (n = 3) or intravenous (IV) bolus at 1 mg/kg (n = 3) with drug compounds formulated in 0.9% saline. Blood samples were taken (EDTA-K2 blood collection tubes) at eight time points over a 24 hr period (T = 5 min, 15 min, 30 min, 1 h, 2 h, 4 h, 8 h, 24 h) for a total of three blood samples per time point. Blood plasma was processed by aliquoting 3 uL of each sample into a 1.5 mL tube. Samples were quenched by addition of a 100 ng/mL internal standard solution in methanol (Labetalol & tolbutamide & Verapamil & dexamethasone & glyburide & Celecoxib, 1:1:1:1:1:1 w/w). The mixture was vortexed and then centrifuged for 15 min at 12,000 x g (4 °C). A 55 uL aliquot of supernatant was transferred to a 96-well plate, centrifuged for 5 min at 3,220 x g (4 °C), then the supernatant was directly injected for LC-MS/MS analysis. Samples were quantified based on a calibration curve of known concentrations. PK parameters were determined by linear/log trapezoidal calculation method using Phoenix WinNonlin 6.3.

### In vivo experiments

All animal studies were carried out according to Stanford University Administrative Panel on Laboratory Animal Care (APLAC) and conducted by Transgenic knockout and Tumor Model Center. 4 to 6 weeks-old male immunodeficient NSG mice (NOD.Cg-Prkdcscid Il2rgtm1Wjl/SzJ) were obtained from The Jackson Laboratory. Mice were IV injected with 1 ×10^6^ NALM6 expressing luciferase (Day 0), 4 days later IV injected with 3 x 10^6^ 2D VCR-CAR+ T cells or mock T cells (Day 4). Beginning on Day 5, select groups received daily IP injections of PBF004 at 10 mg/kg for 21 days after 2D VCR-CAR+ T or mock T cells administration (ending on Day 25). In all experiments, leukemia burden was evaluated 3 times per week using the AMI imager (Spectral Instruments Imaging). Briefly, sedated mice were injected intraperitoneally with 150mg/kg D-luciferin (PerkinElmer), 10 minutes later images were acquired using auto exposure setting and reimaged for 15 and 30 second exposure. Luminescence images (photons/second, p/s) were analyzed using AURA software (Spectral Instruments Imaging). Mice were sacrificed when they began displaying signs of clinical leukemia or luminescence >1010 p/s, spleen was harvested for cell counting and phenotyping for each animal.

## Acknowledgements

The authors thank all members from the Lei Stanley Qi lab for facilitating experiments and useful discussion. The authors thank Dr. Robbie Majzner for providing the Nalm6 cells and useful discussion. The authors thank Nghi Le and Ian Anderson from the Stanford University Protein and Nucleic Acid Facility for DNA origami synthesis. *In vivo* experiments were contracted to the Stanford Transgenic, Knockout and Tumor model Center (TKTC). Chemical synthesis was contracted to WuXi Apptec Limited. The authors thank Nirk E. Quispe Calla, Jasmine Sosa and Hong Zeng from the Stanford Transgenic, Knockout and Tumor model Center (TKTC) for animal experiments. P.B.F acknowledges support by Stanford Maternal & Child Health Research Institute (MCHRI). M.C. acknowledges support by the National Science Foundation Graduate Research Fellowship Program, the ARCS Foundation Scholarship, and the Ruth L. Kirschstein National Research Service Awards for Individual Predoctoral Fellowship (F31AI164936). X.C. acknowledges support by the Stanford Bio-X SIGF Fellowship. L.S.Q. acknowledges support by the Li Ka Shing Foundation, National Science Foundation CAREER award, National Institutes of Health, Stanford Maternal & Child Health Research Institute (MCHRI), and California Institute for Regenerative Medicine (CIRM). This work is supported by National Science Foundation CAREER award (Award #2046650), National Cancer Institute (Grant # 1U01DK127405, 1R01CA266470-01A1), and the Stanford MCHRI through the Uytengsu-Hamilton 22q11 Neuropsychiatry Research Award Program. L.S.Q. is a Chan Zuckerberg Biohub – San Francisco investigator.

## Author Contributions

P.B.F. and L.S.Q. conceived the idea. P.B.F., M.C., L.S.Q. planned the experiments. P.B.F. designed plasmids. P.B.F., D.R., X.C. cloned plasmids. P.B.F., M.C., X.C. performed cell culture experiments. H.W. performed confocal microscopy imaging experiments and analyzed imaging data. P.B.F. designed and synthesized VCLs. J.G. performed chemical synthesis of a biotin dendrimer compound. P.B.F., M.C., and L.S.Q. analyzed the experimental data. P.B.F., M.C., and L.S.Q wrote the manuscript. L.S.Q secured funding. All authors read and commented on the manuscript.

---

**Figure S1.**
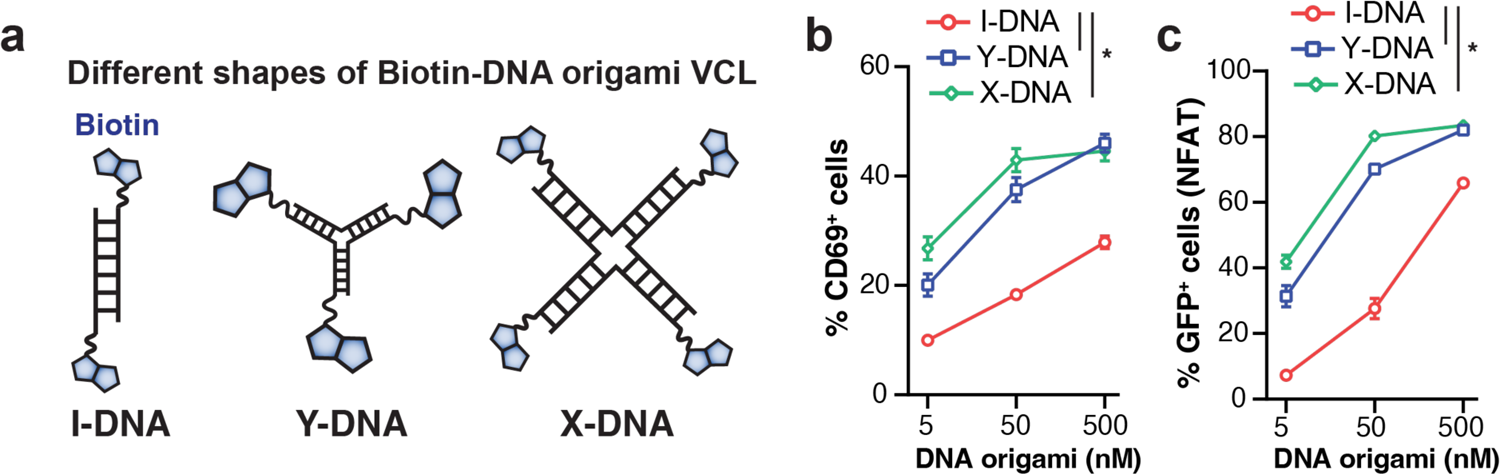
Characterization of VCL valency impact on T cell activation. **a**, Schematic showing multivalent DNA origami based on the I-shape DNA origami (valent=2), Y-shape DNA (valency=3), or X-shape DNA (valency=4), each with ends conjugated with biotin. **b-c**, Characterization of mSA VCR-expressing Jurkat cells, in the presence of various doses of I-(red), Y-(blue), or X-shaped (green) DNA origami for CD69+% T cell activation (**b**) or NFAT signaling activation (**c**). Data contain technical replicates of two biological replicates and are statistically analyzed using a student’s t test, * = p < 0.05.

**Figure S2.**
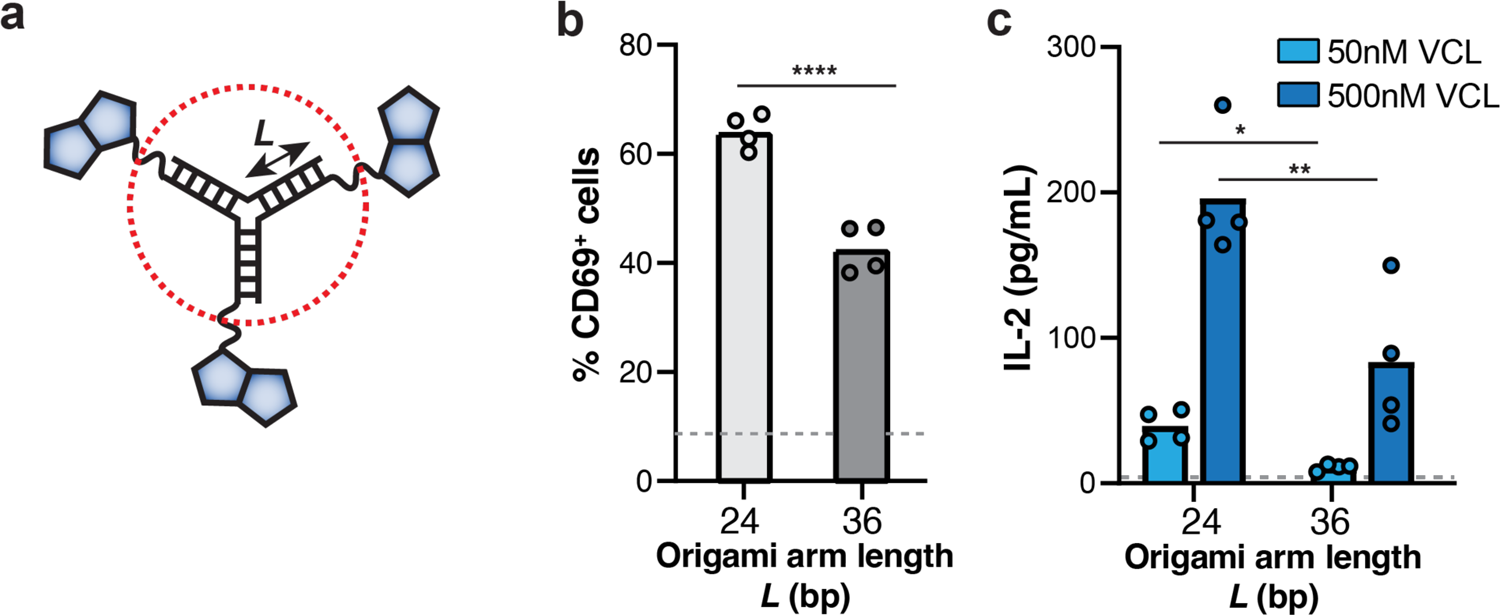
Characterization of receptor proximity impact on T cell activation. **a**, Schematic showing adjusting the arm length of the Y-shape DNA origami VCL (24 vs 36 bp). A small-size VCL presumably induces closer proximity between interacting VCRs. **b-c**, Characterization of mSA VCR-expressing Jurkat cells, in the presence of 24-bp or 36-bp DNA origami VCL for CD69+% T cell activation (**b**) or IL2 production (**c**). Data contain technical replicates of two biological replicates and are statistically analyzed using a student’s t test, * = p < 0.05, ** = p < 0.01, **** = p < 0.0001.

**Figure S3.**
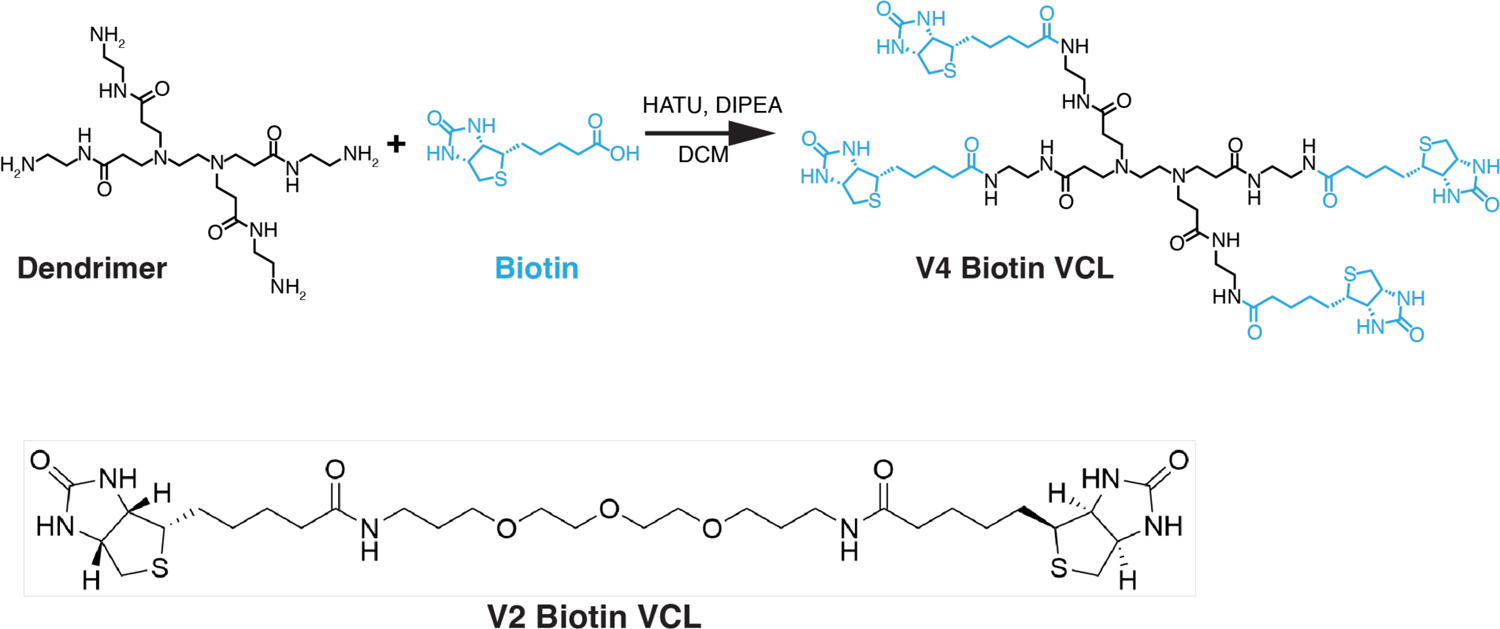
Structures of the dendrimer-based tetravalent V4 Biotin VCL (top) or V2 Biotin VCL (bottom).

**Figure S4.**
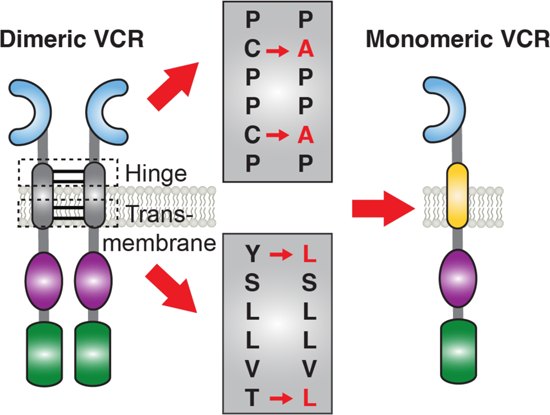
Mutation strategy to create a monomeric mSA VCR. The hinge and transmembrane domains are mutated to create a monomeric version of VCR by disrupting C-C disulfide bonds in the hinge and hydrophobia tyrosine residues in the transmembrane domain.

**Figure S5.**
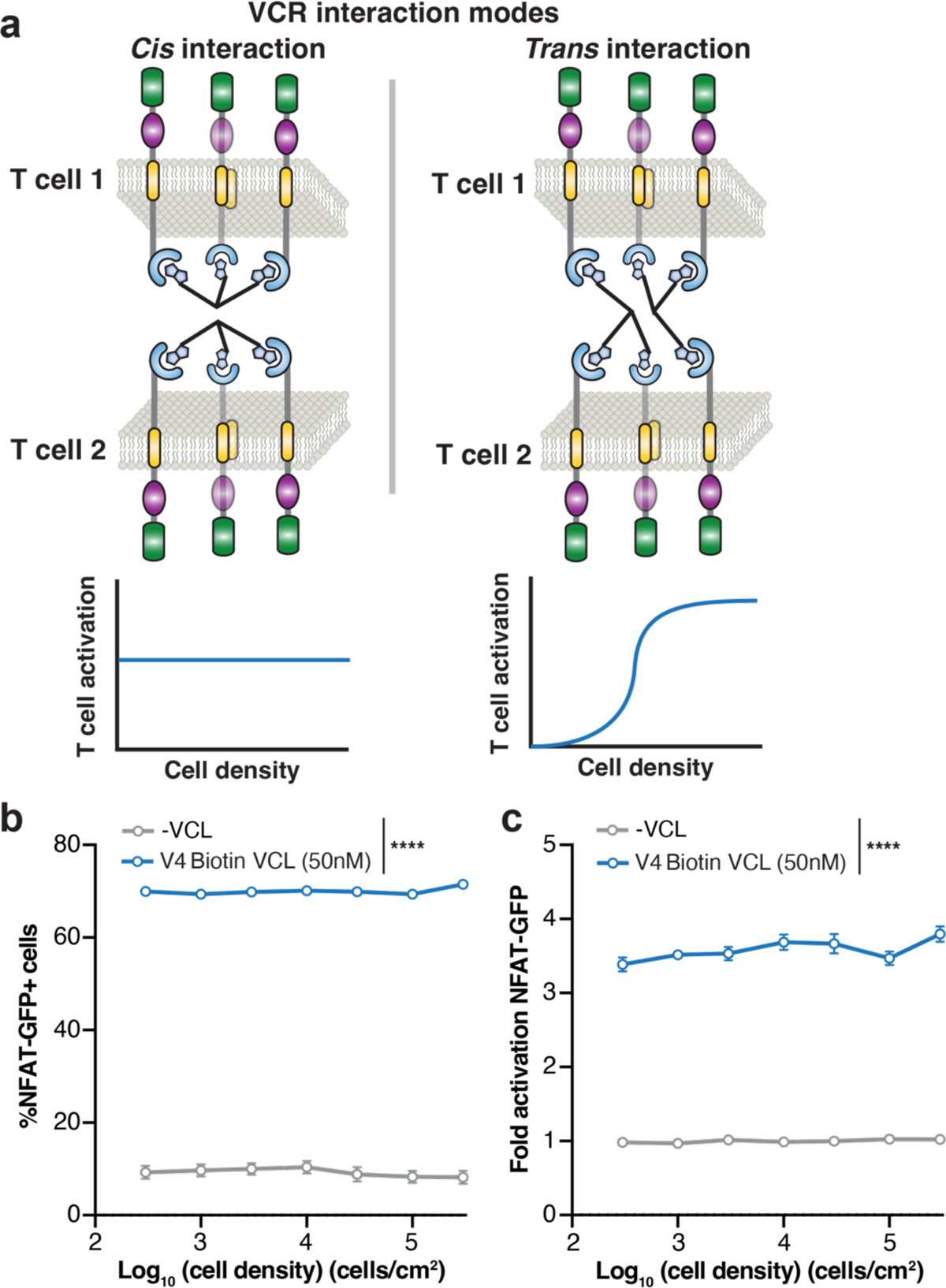
Characterization of cis vs. trans interaction among VCRs for inducing T cell activation. **a**, Schematic showing differences between cis vs. trans interactions among VCRs and how that might lead to different T cell activation responses to cell density. **b-c**, Experimental characterization of mSA VCR Jurkat cells with increasing cell density, without (grey) and with (blue) 50nM V4 Biotin VCL, for NFAT-GFP+% cells (**b**) and fold of activation for NFAT-GFP (**c**). Data are technical replicates of two biological replicates and statistically analyzed using a paired student’s t test, both with **** = p < 0.0001.

**Figure S6.**
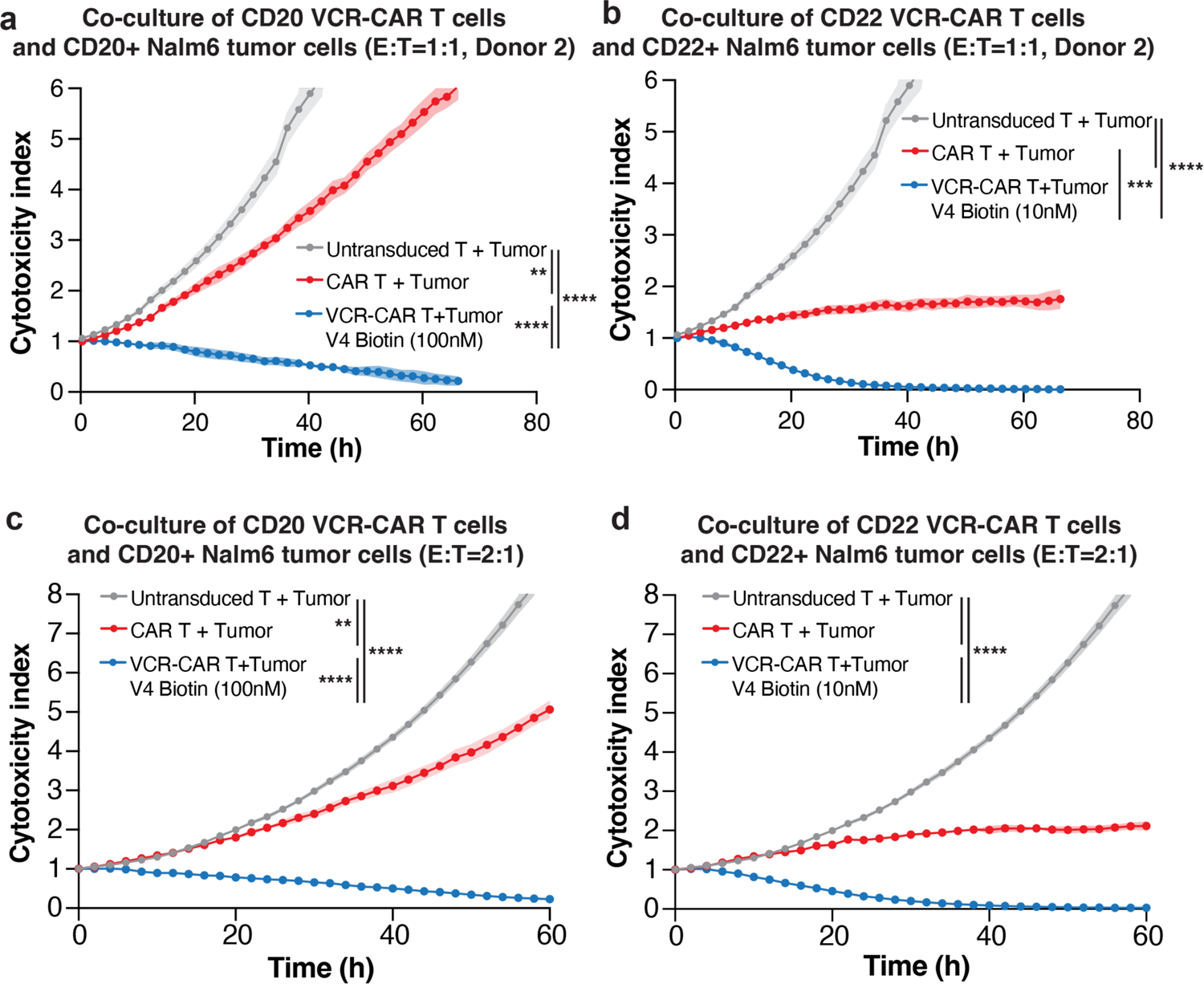
Characterization of VCL-inducible VCR for enhancing CD20 or CD22 CAR activity for tumor killing. **a-d**, all diagrams show real-time live-cell microscopy imaging of the co-culture of antigen-targeting VCR-CAR primary human T cells with antigen^+^ Nalm6 tumor cells in the presence of V4 Biotin VCL (blue), compared to untransduced T + tumor (grey) and CAR T + tumor co-culture (red) samples. **a**, CD20-targeting, E:T=1:1, donor 2; **b**, CD22-targeting, E:T=1:1, donor 2; **c**, CD20-targeting, E:T=2:1; **d**, CD22-targeting, E:T=2:1. Each graph represents one donor with technical triplicates as mean and standard deviation. Data are statistically analyzed using repeated measures ANOVA, ** = p < 0.01, *** = p < 0.001, **** = p < 0.0001.

**Figure S7.**
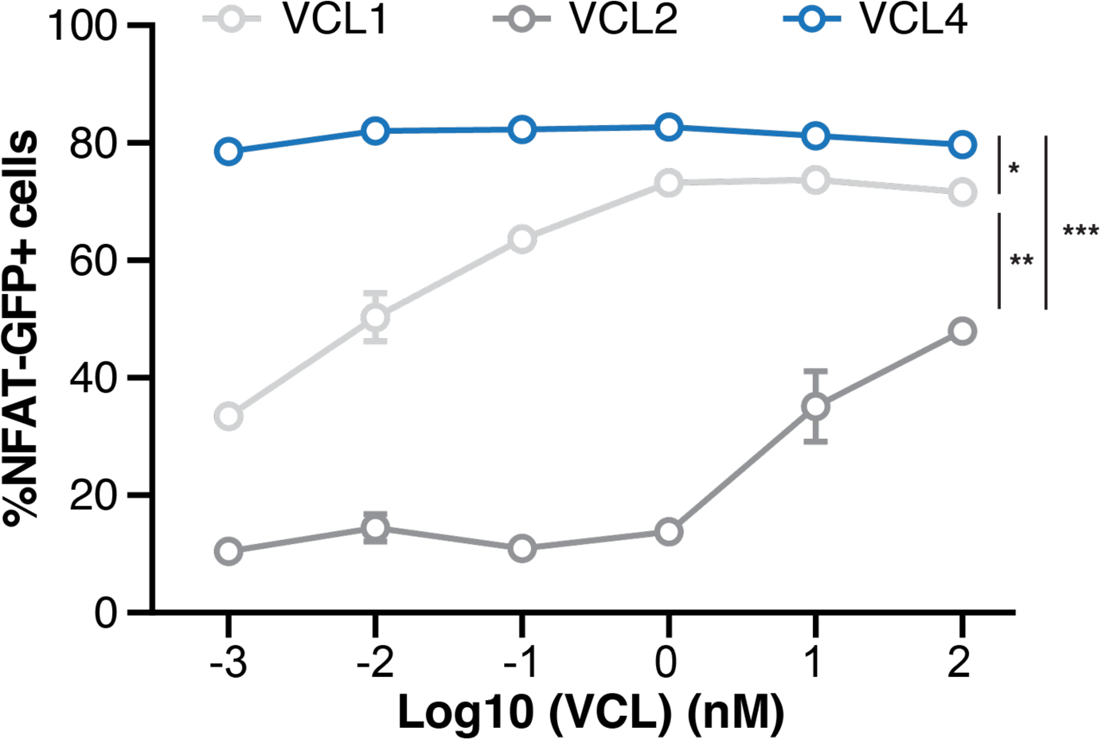
Characterization of GFP+% cells with activated NFAT signaling in Jurkat cells at various drug concentrations of lead VCL compounds (VCL1, VCL2, VCL4). Data contain technical duplicates of two biological replicates and are statistically analyzed using a paired student’s t test, * = p < 0.05, ** = p < 0.01, *** = p < 0.001.

**Figure S8.**
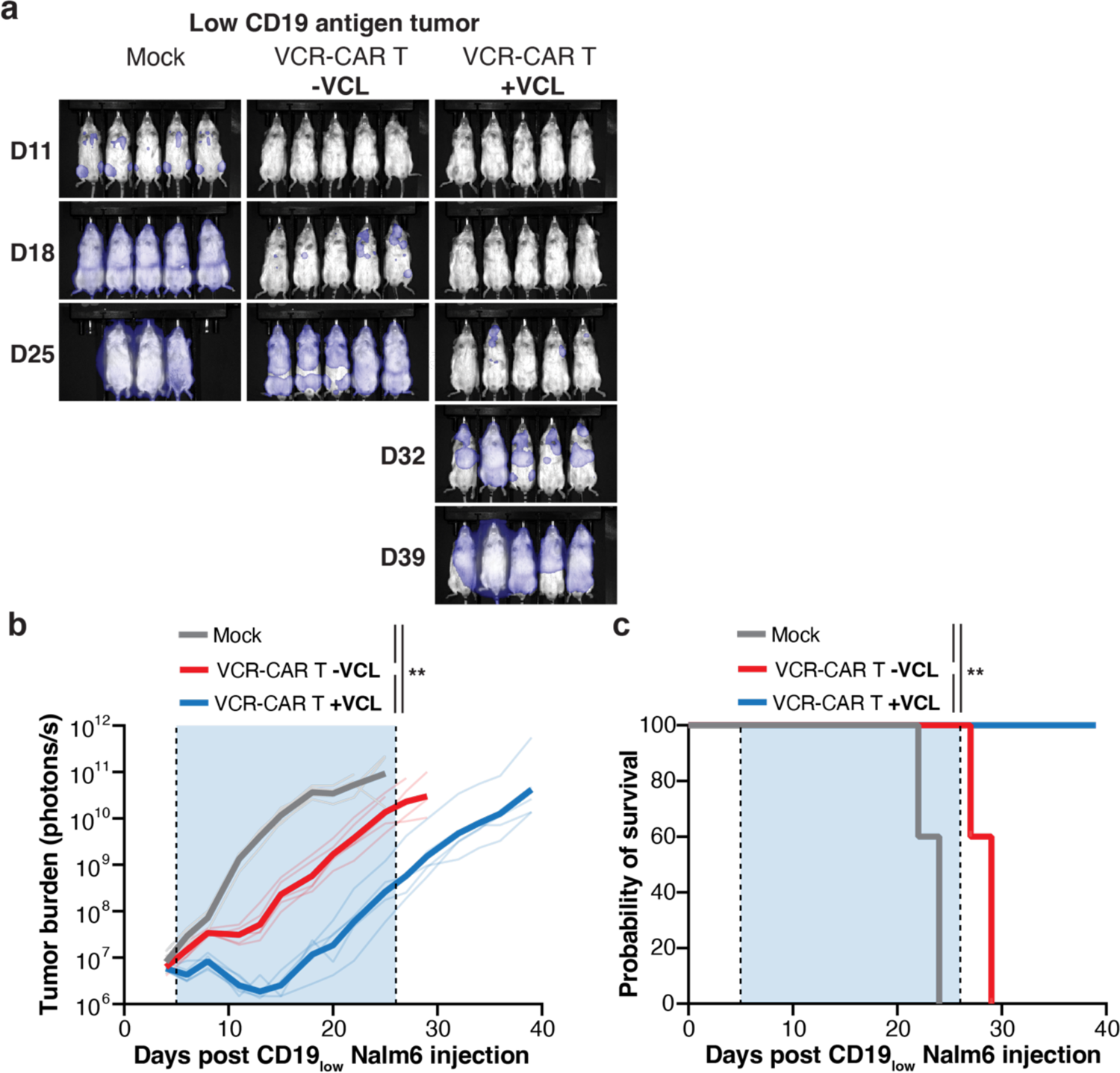
Characterization of using multivalent VCL drug to VCR-CAR T cells for killing low-antigen tumor in vivo. **a**, BLI images showing groups of mice injected with mock T cells, VCR-CAR (CD19 scFv-mSA-4-1BB-CD3ζ) T cells without or with VCL. All mice were injected with CD19_low_ Nalm6 cells in the same experimental design as **Fig. 5f**. **b**, Characterization of CD19_low_ Nalm6 tumor burden (photon/s) for VCL-dosed mice carrying VCR-CAR T cells without VCL (red) or with VCL (blue), and mock (grey). Data are statistically analyzed using repeated measure ANOVA, ** = p < 0.01. **c**, Viability curve for VCL-dosed mice carrying VCR-CAR T cells without VCL (red) or with VCL (blue), and mock (grey). Data from 5 mice are analyzed using a log rank test, ** p < 0.01.

